# Computational Prediction of the Comprehensive SARS-CoV-2 vs. Human Interactome to Guide the Design of Therapeutics

**DOI:** 10.1101/2020.03.29.014381

**Authors:** Kevin Dick, Kyle K. Biggar, James R. Green

## Abstract

Understanding the disease pathogenesis of the novel coronavirus, denoted SARS-CoV-2, is critical to the development of anti-SARS-CoV-2 therapeutics. The global propagation of the viral disease, denoted COVID-19 (“coronavirus disease 2019”), has unified the scientific community in searching for possible inhibitory small molecules or polypeptides. Given the known interaction between the human ACE2 (“Angiotensin-converting enzyme 2”) protein and the SARS-CoV virus (responsible for the coronavirus outbreak *circa*. 2003), considerable focus has been directed towards the putative interaction between the SARS-CoV-2 Spike protein and ACE2. However, a more holistic understanding of the SARS-CoV-2 vs. human inter-species interactome promises additional putative protein-protein interactions (PPI) that may be considered targets for the development of inhibitory therapeutics.

To that end, we leverage two state-of-the-art, sequence-based PPI predictors (PIPE4 & SPRINT) capable of generating the comprehensive SARS-CoV-2 vs. human interactome, comprising approximately 285,000 pairwise predictions. Of these, we identify the high-scoring subset of human proteins predicted to interact with each of the 14 SARS-CoV-2 proteins by both methods, comprising 279 high-confidence putative interactions involving 225 human proteins. Notably, the Spike-ACE2 interaction was the highest ranked for both the PIPE4 and SPRINT predictors, corroborating existing evidence for this PPI. Furthermore, the PIPE-Sites algorithm was used to predict the putative subsequence that might mediate each interaction and thereby inform the design of inhibitory polypeptides intended to disrupt the corresponding host-pathogen interactions.

We hereby publicly release the comprehensive set of PPI predictions and their corresponding PIPE-Sites landscapes in the following DataVerse repository: 10.5683/SP2/JZ77XA. All data and metadata are released under a CC-BY 4.0 licence. The information provided represents theoretical modeling only and caution should be exercised in its use. It is intended as a resource for the scientific community at large in furthering our understanding of SARS-CoV-2.

## 1 Introduction

The novel coronavirus pandemic has galvanized the research community into the investigation of the SARS-CoV-2 virus and the COVID-19 disease it manifests in humans [1]. Research has progressed with unprecedented speed due, in large part, to the rapid determination of the SARS-CoV-2 genome and proteome. These data enable the research community to collectively contribute to the study and understanding of SARS-CoV-2 and its disease pathogenesis.

Promisingly, many computational approaches have been rapidly deployed to increase our understanding of SARS-CoV-2, including protein function, three-dimensional (3D) protein structures, and possible target regions for small inhibitory molecules [2, 3]. Through the use of publication preprint platforms, this information can be immediately disseminated, albeit, with the disclaimer of “non-peer-reviewed” research. Two notable examples include the use of DeepMind’s recently published AplhaFold protein structure predictor [2] to predict the 3D protein structure of each of the SARS-CoV-2 proteins, and the use of the SUMMIT high-performance computing (HPC) infrastructure to perform large-scale virtual docking simulations as a form of high-throughput screening to identify small inhibitory molecules [3]. Given that the Spike protein from the original SARS coronavirus, SARS-CoV, is known to interact with the human Angiotensin-Converting Enzyme 2 (ACE2), current efforts are focused to better characterize the SARS-CoV-2 Spike protein and its putative interaction with the ACE2 protein.

Similar efforts are being made to understand the functional and evolutionary characteristics of the SARS-CoV-2 proteome, including the determination of evolutionary conserved functional regions between related viruses to inform the use of anti-viral therapeutics [4]. Given the unique infectivity characteristics of this novel coronavirus, the need for effective anti-viral therapeutics is pressing. The long viral incubation period, during which an individual is simultaneously contagious and asymptomatic, has resulted in rapid global proliferation. Leveraging what is known from the original SARS-CoV outbreak, *circa*. 2003, and related viral families, this work contributes predicted protein-protein interaction (PPI) networks to guide researchers and form the basis of testable wet laboratory hypotheses.

Coronaviruses share many similarities to the influenza viruses in that they are both enveloped, single-stranded, and helical RNA-viruses among the Group IV viral families [5]. The four coronaviruses known to commonly infect humans are believed to have evolved such that they maximize proliferation within a population. This evolved strategy involves sickening, but not ultimately killing, their hosts. By contrast, the two prior novel coronavirus outbreaks—SARS (severe acute respiratory syndrome) and MERS (Middle East respiratory syndrome, named for where the first outbreak occurred)—arose in humans after cross-species jumps from animals, as was H5N1 (the avian influenza). These latter diseases were highly fatal to humans, with a few mild or asymptomatic cases. A greater proportion of mild or asymptomatic cases would have resulted in wide-spread disease, however, SARS and MERS each ultimately killed fewer than 1,000 people.

The scientific community has been spurred into action in response to the recent COVID-19 outbreak, building on decades of basic research characterising this virus family. Labs at the forefront of the outbreak response shared genomes of the virus in open access databases, which enabled researchers to rapidly develop tests for this novel pathogen. Other labs have shared experimentally-determined and computationally-predicted structures of some of the viral proteins, and still others have shared epidemiological data. We hope to contribute to the scientific effort using the latest version of our sequence-based protein-protein interaction (PPI) predictor, PIPE4 [6] in combination with another state-of-the-art PPI predictor, denoted Scoring PRotein INTeractions (SPRINT) [7]. Finally, the PIPE-Sites algorithm was used to predict the sub-sequence regions with a high likelihood of mediating the physical interaction between two given pairs[8].

The rapidity of our response is thanks, in part, to having produced an analogous study during the emergence of the Zika Virus outbreak of 2015, where our sequence-based PPI prediction method (PIPE) was used to identify putative human-Zika inter-species PPIs and inform possible synthetic biology approaches for novel interventions and therapeutics [9]. In the present study, of the ∼ 285,000 predicted pairs, we selected only the highly conservative set of predicted interactions for each of the 14 SARS-CoV-2 proteins resulting in the identification of 225 putative human protein targets. We publicly released these predictions and related meta-data for use by the broader scientific community in the following DataVerse repository: 10.5683/SP2/JZ77XA, [10].

## 2 Methods

Following from a previous study of the Zika virus in 2016 [9], we defined two prediction schemas from which to train the PPI predictors. First, the *all* schema, contains the maximum available number of known virus-host PPIs regardless of the evolutionary distance between those viruses and the target virus (*i*.*e*. SARS-CoV-2). This schema groups all viruses into a “viral” collection to serve as a proxy for SARS-CoV-2. The second schema, denoted *proximal*, is a subset of the *all* schema, where only the PPIs from evolutionarily related organisms are considered. In both schemas, to avoid possible overfitting, the previously known SARS-CoV Spike vs. ACE2 interaction was removed. We retained the other four known interactions between SARS-CoV and human. At present, only the results of the *all* schema are presented and analyzed. A subsequent version of this article will additionally provide the results of the *proximal* schema.

The dataset of experimentally elucidated human-virus PPIs was obtained from the VirusMentha database [11]. These 10,693 known PPIs are used to train the PPI predictors and infer new putative interactions between human proteins and the SARS-CoV-2 proteome. For the *all* schema, the proteomes of the 43 viral families were collected from Uniprot and are summarized in the appendix Table 4. In anticipation of the generation of the predicted interactome using the *proximal* schema, we tabulate the 689 training PPI and the Group IV viral families over which they are distributed (Table 1). Finally, the human reference proteome (UP000005640) was obtained from Uniprot, retaining only the high-quality “Reviewed” Swiss-Prot proteins.

**Table 1:**
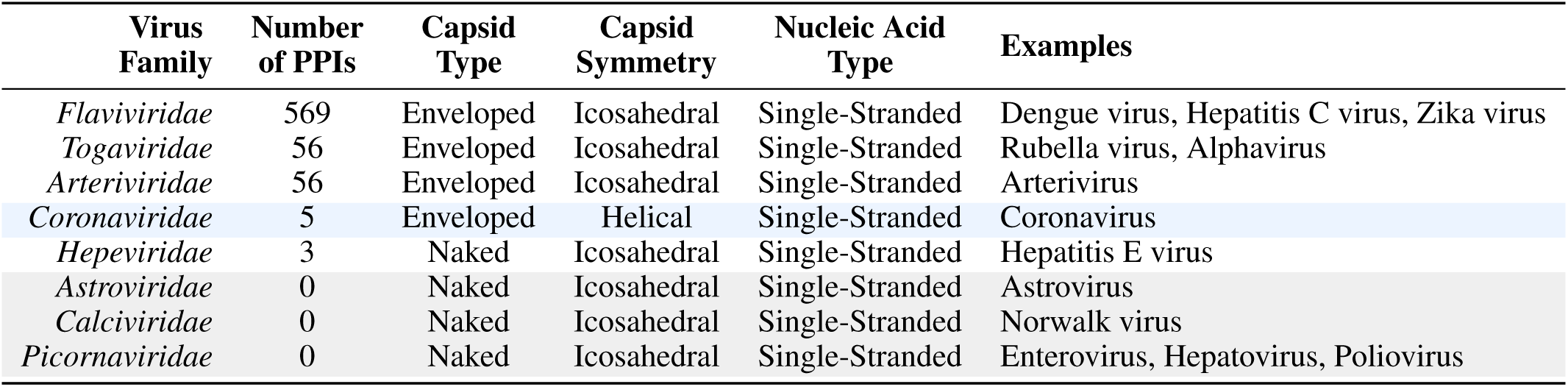
Group IV Viral Families and their Number of PPIs used in the Proximal Prediction Schema.

**Table 2:** The 14 Proteins in the SARS-CoV-2 Proteome.

| Uniprot Acc. | Gene Name | Protein Name | Protein Function |
| --- | --- | --- | --- |
| P0DTD1 | R1A_WCPV | Replicase polyprotein 1a (R1a) | Viral transcription/replication |
| P0DTC1 | R1AB_WCPV | Replicase polyprotein 1ab (R1ab) | Viral transcription/replication, ribosomal frameshift |
| P0DTC2 | SPIKE_WCPV | Spike glycoprotein (S) | Attachment and entry |
| P0DTC3 | AP3A_WCPV | Protein 3a | ESCRT-independent budding |
| P0DTC4 | VEMP_WCPV | Envelope small membrane protein (E) | ESCRT-independent budding |
| P0DTC5 | VME1_WCPV | Membrane protein (M) | Virion morphogenesis |
| P0DTC6 | NS6_WCPV | Non-structural protein 6 | Unknown; possibly host-virus modulation |
| P0DTC7 | NS7A_WCPV | Protein 7a (NS7A) | Unknown; possibly host-virus modulation |
| P0DTD8 | NS7B_WCPV | Protein 7b (NS7B) | Unknown; possibly host-virus modulation |
| P0DTC8 | NS8_WCPV | Non-structural protein 8 (NS8) | Unknown; possibly host-virus modulation |
| P0DTC9 | NCAP_WCPV | Nucleoprotein (N) | Viral genome packaging |
| P0DTD3 | Y14_WCPV | Uncharacterized protein 14 | Unknown; possibly host-virus modulation |
| P0DTD2 | ORF9B_WCPV | Protein 9b | Unknown; possibly host-virus modulation |
| A0A663DJA2 | A0A663DJA2_9BETC | Hypothetical ORF10 protein | Presumably not expressed |

**Table 3:**
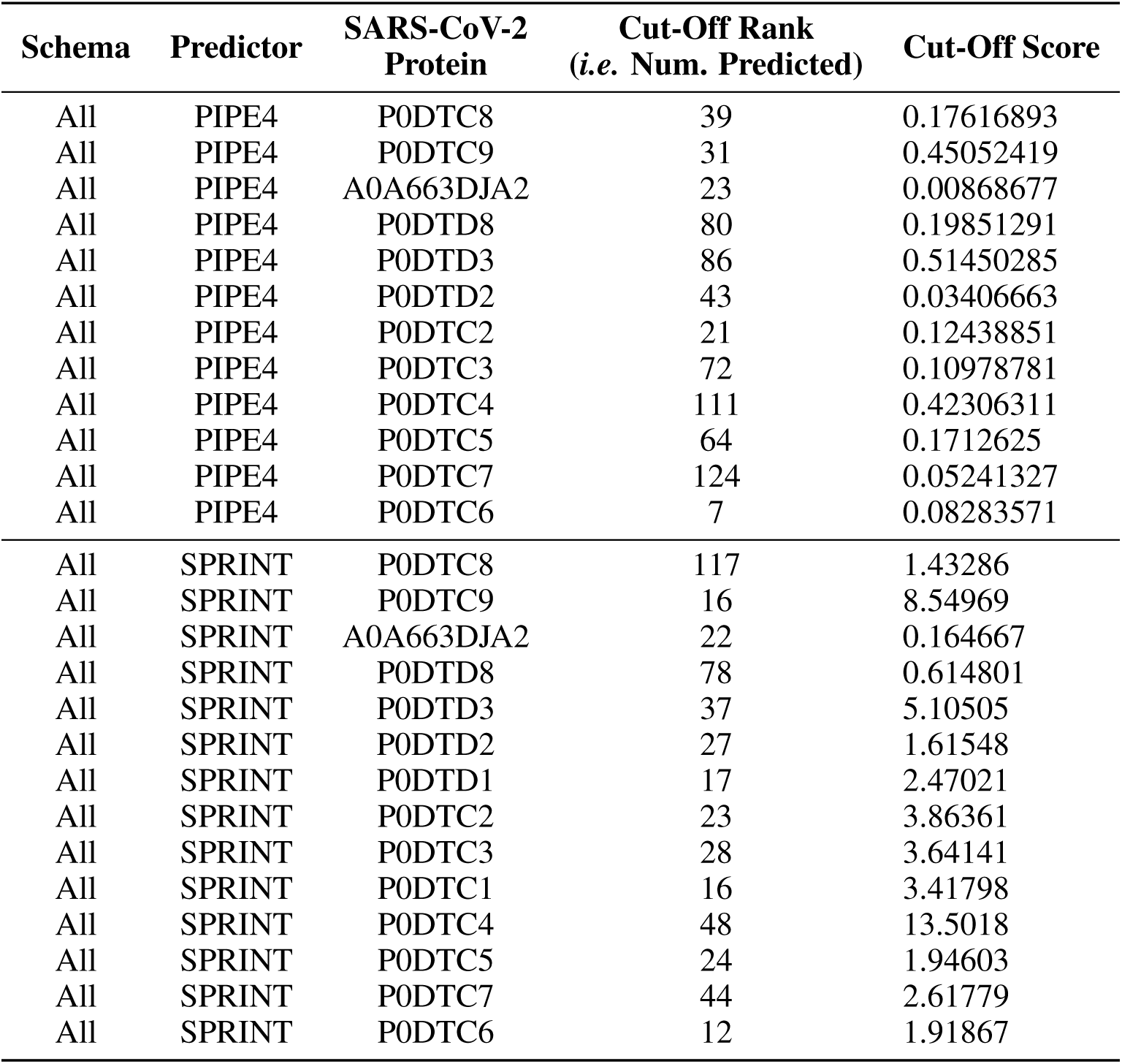
Summary of the Number of Predicted Interactions.

| Schema | Predictor | SARS-CoV-2 Protein | Cut-Off Rank<br>(i.e. Num. Predicted) | Cut-Off Score |
| --- | --- | --- | --- | --- |
| All | PIPE4 | P0DTC8 | 39 | 0.17616893 |
| All | PIPE4 | P0DTC9 | 31 | 0.45052419 |
| All | PIPE4 | A0A663DJA2 | 23 | 0.00868677 |
| All | PIPE4 | P0DTD8 | 80 | 0.19851291 |
| All | PIPE4 | P0DTD3 | 86 | 0.51450285 |
| All | PIPE4 | P0DTD2 | 43 | 0.03406663 |
| All | PIPE4 | P0DTC2 | 21 | 0.12438851 |
| All | PIPE4 | P0DTC3 | 72 | 0.10978781 |
| All | PIPE4 | P0DTC4 | 111 | 0.42306311 |
| All | PIPE4 | P0DTC5 | 64 | 0.1712625 |
| All | PIPE4 | P0DTC7 | 124 | 0.05241327 |
| All | PIPE4 | P0DTC6 | 7 | 0.08283571 |
| All | SPRINT | P0DTC8 | 117 | 1.43286 |
| All | SPRINT | P0DTC9 | 16 | 8.54969 |
| All | SPRINT | A0A663DJA2 | 22 | 0.164667 |
| All | SPRINT | P0DTD8 | 78 | 0.614801 |
| All | SPRINT | P0DTD3 | 37 | 5.10505 |
| All | SPRINT | P0DTD2 | 27 | 1.61548 |
| All | SPRINT | P0DTD1 | 17 | 2.47021 |
| All | SPRINT | P0DTC2 | 23 | 3.86361 |
| All | SPRINT | P0DTC3 | 28 | 3.64141 |
| All | SPRINT | P0DTC1 | 16 | 3.41798 |
| All | SPRINT | P0DTC4 | 48 | 13.5018 |
| All | SPRINT | P0DTC5 | 24 | 1.94603 |
| All | SPRINT | P0DTC7 | 44 | 2.61779 |
| All | SPRINT | P0DTC6 | 12 | 1.91867 |

**Table 4:** Proteomes of the Majority of Organisms Considered in the *All* Schema.

| Organism | Taxonomy Id | Proteome Acc. |
| --- | --- | --- |
| Rotavirus A | 9913 | UP000106064 |
| Sindbis virus (SINV) | 11034 | UP000006710 |
| Rubella virus (strain M33) (RUBV) | 11043 | UP000007143 |
| Dengue virus 1 | 11053 | UP000101782 |
| Dengue virus 2 | 11060 | UP000096836 |
| Dengue virus type 2 (strain Thailand/NGS-C/1944) (DENV-2) | 11065 | UP000007196 |
| Japanese encephalitis virus | 11072 | UP000121923 |
| Kunjin virus | 11077 | UP000100779 |
| Kunjin virus (strain MRM61C) | 11078 | UP000099558 |
| West Nile virus (WNV) | 11082 | UP000102709 |
| Tick-borne encephalitis virus | 11084 | UP000140821 |
| Classical swine fever virus | 11096 | UP000106488 |
| Bovine viral diarrhea virus (BVDV) (Mucosal disease virus) | 11099 | UP000155116 |
| Hepatitis C virus genotype 1a (isolate H) (HCV) | 11103 | UP000000518 |
| Hepatitis C virus genotype 1a (isolate 1) (HCV) | 11104 | UP000008855 |
| Hepatitis C virus genotype 1b (isolate BK) (HCV) | 11105 | UP000007413 |
| Hepatitis C virus genotype 1a (isolate H) (HCV) | 11108 | UP000000518 |
| Hepatitis C virus genotype 2a (isolate HC-J6) (HCV) | 11113 | UP000002682 |
| Hepatitis C virus genotype 1b (isolate Japanese) (HCV) | 11116 | UP000008095 |
| Human coronavirus 229E (HCoV-229E) | 11137 | UP000006716 |
| Hepatitis E virus (HEV) | 12461 | UP000106507 |
| Porcine reproductive and respiratory syndrome virus (PRRSV) | 28344 | UP000146080 |
| Dengue virus type 2 (strain Thailand/16681/1984) (DENV-2) | 31634 | UP000180751 |
| Dengue virus type 2 (strain 16681-PDK53) (DENV-2) | 31635 | UP000008390 |
| Hepatitis C virus genotype 1b (isolate Taiwan) (HCV) | 31645 | UP000002679 |
| Hepatitis E virus genotype 1 (isolate Human/Burma) (HEV-1) | 31767 | UP000007243 |
| Hepatitis E virus genotype 2 (isolate Human/Mexico) (HEV-2) | 31768 | UP000007245 |
| Bovine viral diarrhea virus 2 | 54315 | UP000129869 |
| Alkhurma hemorrhagic fever virus (ALKV) | 172148 | UP000097483 |
| Human SARS coronavirus (SARS-CoV) | 227859 | UP000000354 |
| Porcine epidemic diarrhea virus (strain CV777) (PEDV) | 229032 | UP000008159 |
| SARS coronavirus Frankfurt 1 | 229992 | UP000113286 |
| Porcine torovirus | 237020 | UP000269215 |
| Human coronavirus NL63 (HCoV-NL63) | 277944 | UP000103541 |
| Hepatitis C virus genotype 1b (isolate Con1) (HCV) | 333284 | UP000007414 |
| Hepatitis C virus genotype 2a (isolate JFH-1) (HCV) | 356411 | UP000008096 |
| Breda virus 1 (BRV-1) | 360393 | UP000000355 |
| Dengue virus type 4 (strain Dominica/814669/1981) (DENV-4) | 408871 | UP000108177 |
| Hepatitis C virus genotype 1b (strain HC-J4) (HCV) | 420174 | UP000008094 |
| Hepatitis C virus genotype 1b (isolate HC-J1) (HCV) | 421877 | UP000008093 |
| Hepatitis C virus genotype 1b (isolate HCR6) (HCV) | 421879 | UP000008100 |
| Hepatitis E virus genotype 4 (isolate Human/China/T1) (HEV-4) | 509627 | UP000007242 |
| Hepatitis E virus genotype 1 (isolate Human/India/Hyderabad) | 512346 | UP000007244 |

### 2.1 The SARS-CoV-2 Proteome

The proteome of SARS-CoV-2 was obtained from the Uniprot pre-release available at SARS-CoV-2 Pre-Release, [12], with the disclaimer that these data will become part of a future UniProt release and may be subject to further changes. The 14 SARS-CoV-2 proteins and their function are tabulated in Table 2. Notably, the P0DTC2 Spike glycoprotein is of special interest to this and related work, since its SARS-CoV equivalent is known to interact with the human ACE2 protein.

### 2.2 Computational Protein-Protein Interaction Predictors

The computational prediction of PPIs is a diverse field which encompasses multiple paradigms (*e*.*g*. sequence-, structure-, evolution-, and network-based methods). The shortcomings of one approach are often the strength of another and certain paradigms can be useful in generating insightful interaction interface information. Here, we will discuss the two paradigms with specific relevance to the SARS-CoV-2 pandemic given the current focus of the research community in an effort to develop therapeutics that might slow the progression and impact of COVID-19.

Structure-based methods require knowledge the 3D structure of each of the proteins from the set of known PPIs and also for each of the proteins for which one wishes to make inferences. Consequently, these methods suffer from low coverage throughout a complete proteome and are generally unsuitable for comprehensive interactome predictions. Furthermore, many structure-based methods rely on *de novo* or template-based modelling, which tend to be computationally taxing. Promisingly, the DeepMind team that developed the AlphaFold computation protein structure predictor have publicly released their predictions of the 14 proteins in the SARS-CoV-2 proteome for use by the scientific community, enabling the use of structure-based prediction methods. However, high quality structures are not available for all human proteins and, even with complete 3D structural information of each protein in both organisms’ proteomes, the computational time complexity to elucidate all possible inter-species pairings make these methods prohibitive beyond modestly sized networks. Promisingly, these methods are highly complimentary to other prediction paradigms and can be applied following the initial screening using other, more computationally efficient and high-throughput PPI prediction methods.

At the other computational extreme, sequence-based predictors rely solely upon primary sequence data making them amenable to the investigation of proteome-wide networks. Furthermore, these methods tend to be highly efficient, where individual PPIs can be predicted in a fraction of a second.

#### 2.2.1 The Protein-Protein Interaction Prediction Engine (PIPE4)

PIPE is a sequence-based method of PPI prediction that operates by examining sequence windows on each of the query proteins. If the pair of sequence windows shares significant similarity with a pair of proteins previously known to interact, then evidence for the putative PPI is increased. A similarity-weighted (SW) scoring function uses normalization to account for frequently occurring sequences, not related to PPIs. Given sufficient evidence, a PPI is predicted. PIPE has previously been validated on numerous species for both intra-species and inter-species PPI prediction tasks [13, 14, 15]. Furthermore, the distribution of evidence along the length of each query protein forms a 2D landscape that can indicate the site of interaction (see “Predicted PPI Site of Interaction” subsection 2.2.4 below) [8].

The fourth version of the Protein-protein Interaction Prediction Engine (PIPE4) was recently adapted to improve predictive performance for understudied organisms [6]. That is, species for which the proteome is known, however, the the number of experimentally validated PPIs involving the proteins of this organism is insufficient to train a model to generate the comprehensive interactome. To circumvent this, the PIPE4 algorithm leverages the known PPIs of evolutionarily similar and well-studied organism, serving as a *proxy* training set. Using an approach denoted as *cross-species* PPI prediction, the experimentally validated PPIs from the proxy species are used to train the PPI predictor which is then applied to the proteome of the understudied target organism. Due to the limited availability of known SARS-CoV-2 PPIs, we here use the PPIs from a collection of well-studied and evolutionarily similar proxy viruses to generate these cross-species predictions as depicted in Figure 1.

**Figure 1:**
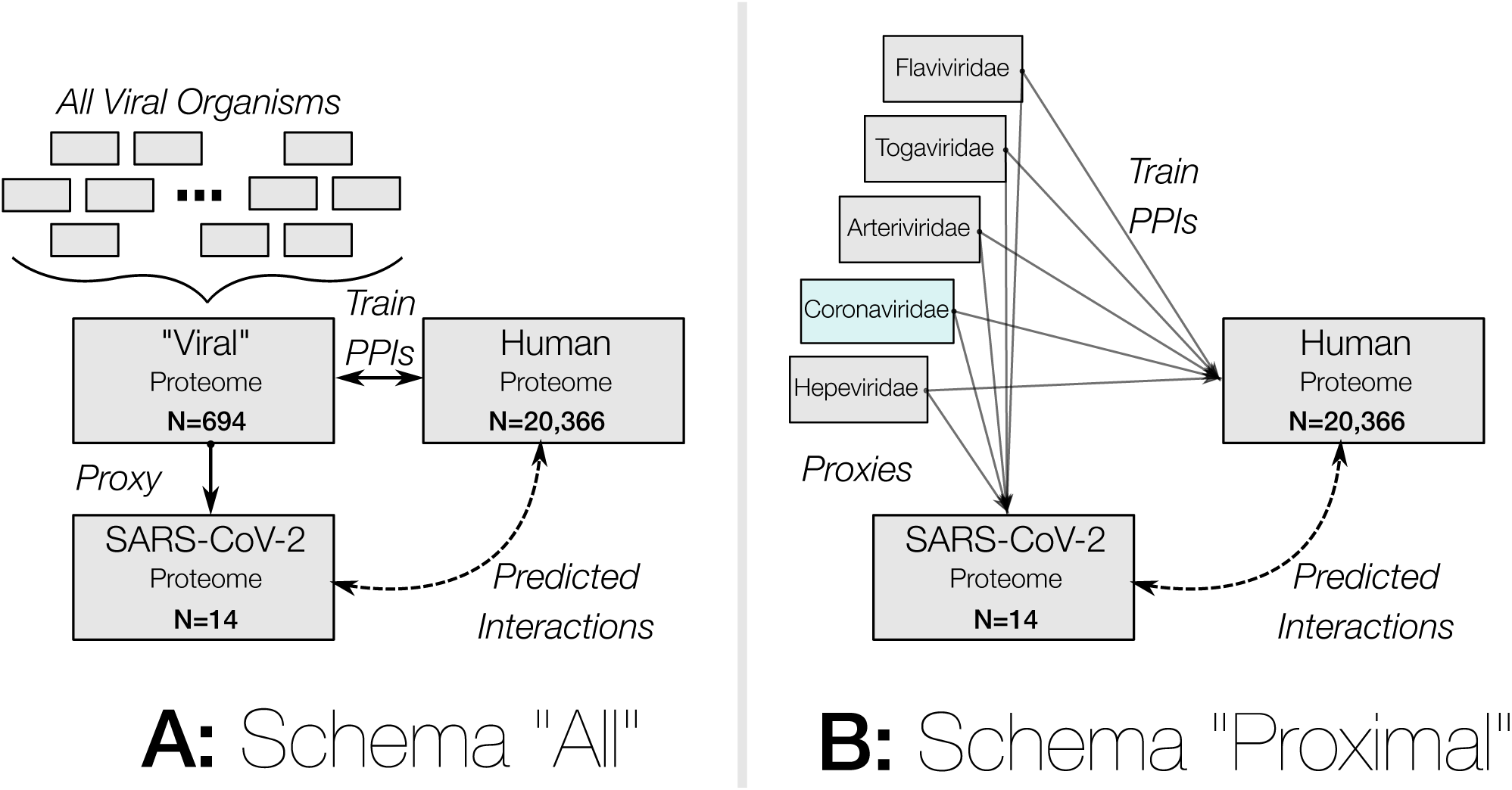
Overview of the Two Prediction Strategies to Generate the SARS-CoV-2 vs. Human Interactome. At present, only the results of the *All* schema are presented in this work.

The PIPE4 algorithm is particularly well-suited to cross- and inter-species PPI prediction schemas, given that the SW-scoring function appropriately normalizes the prevalence of sequence windows within each training and target species proteome [6].

#### 2.2.2 Scoring PRotein INTeractions (SPRINT)

The SPRINT predictor is conceptually similar to PIPE; SPRINT aggregates evidence from previously known PPI interactions, depending on window similarity with the query protein pair, to inform its prediction scores [7]. SPRINT leverages a *spaced seed* approach for determining protein window sequence similarity, where each position in the two windows must either be identical, or do not matter, as defined by the match/don’t_care bits of the spaced seeds. Furthermore, protein sequences are encoded using five bits per amino, enabling the use of highly efficient (SIMD) bitwise operations to rapidly compute protein window similarities and, thereby, score predictions [7]. Unfortunately, the present version of the SPRINT algorithm is not designed to explicitly handle inter- and cross-species prediction, nor to predict the specific subsequence site of interaction between a given pair of proteins. Nonetheless, it is among the only PPI predictors capable of predicting comprehensive interactomes in a timely manner and was demonstrated to outperform other PPI predictors, including the PIPE2 algorithm [7].

#### 2.2.3 Determining an Appropriate Per-Protein Decision Threshold

For each of the 14 SARS-CoV-2 proteins, we predicted their interaction with each of the 20,366 human proteins resulting in 285,124 unique predictions from each of the two predictors considered. From these predicted interactomes, we can plot the rank-ordered distribution of the putative interaction scores involving each of the *single* SARS-CoV-2 proteins separately. This presents an opportunity to develop protein-specific local decision thresholds, where only those interactions scoring significantly above baseline are reported. These one-to-all score curves are based on the underlying assumption that we expect true SARS-CoV-2 vs. human PPIs to be rare, such that the vast majority of prediction scores should fall below the decision threshold. Furthermore, by also plotting the one-to-all curves for each human protein, we can apply the same local decision logic to the reciprocal perspective (while not performed here, this analysis forms the basis of the Reciprocal Perspective method) [16].

Thus, for each one-to-all score curve, a score threshold delineating the “high-scoring” PPIs from the baseline was identified and used to determine the high-confidence interactions. In the absence of known PPIs between SARS-CoV-2 and human, it is difficult to determine a suitable global decision threshold. By instead examining the morphology of the one-to-all score curves for both perspectives, we can qualitatively identify high-scoring pairs. This process can be further automated through the identification of the baseline/knee for each view under the assumption that true PPIs are rare and high-scoring, while non-interacting pairs tend to generate scores residing below the knee in the baseline. In Figure 2, we overlay the one-to-all score curves for each SARS-CoV-2 protein and “zoom’ into the high-score/low-rank region to emphasize that the selection of a single global top-*k* or score threshold would inappropriately exclude relatively high-scoring pairs within specific SARS-CoV-2 proteins, while admitting too many low-scoring putative PPI for other proteins.

**Figure 2:**
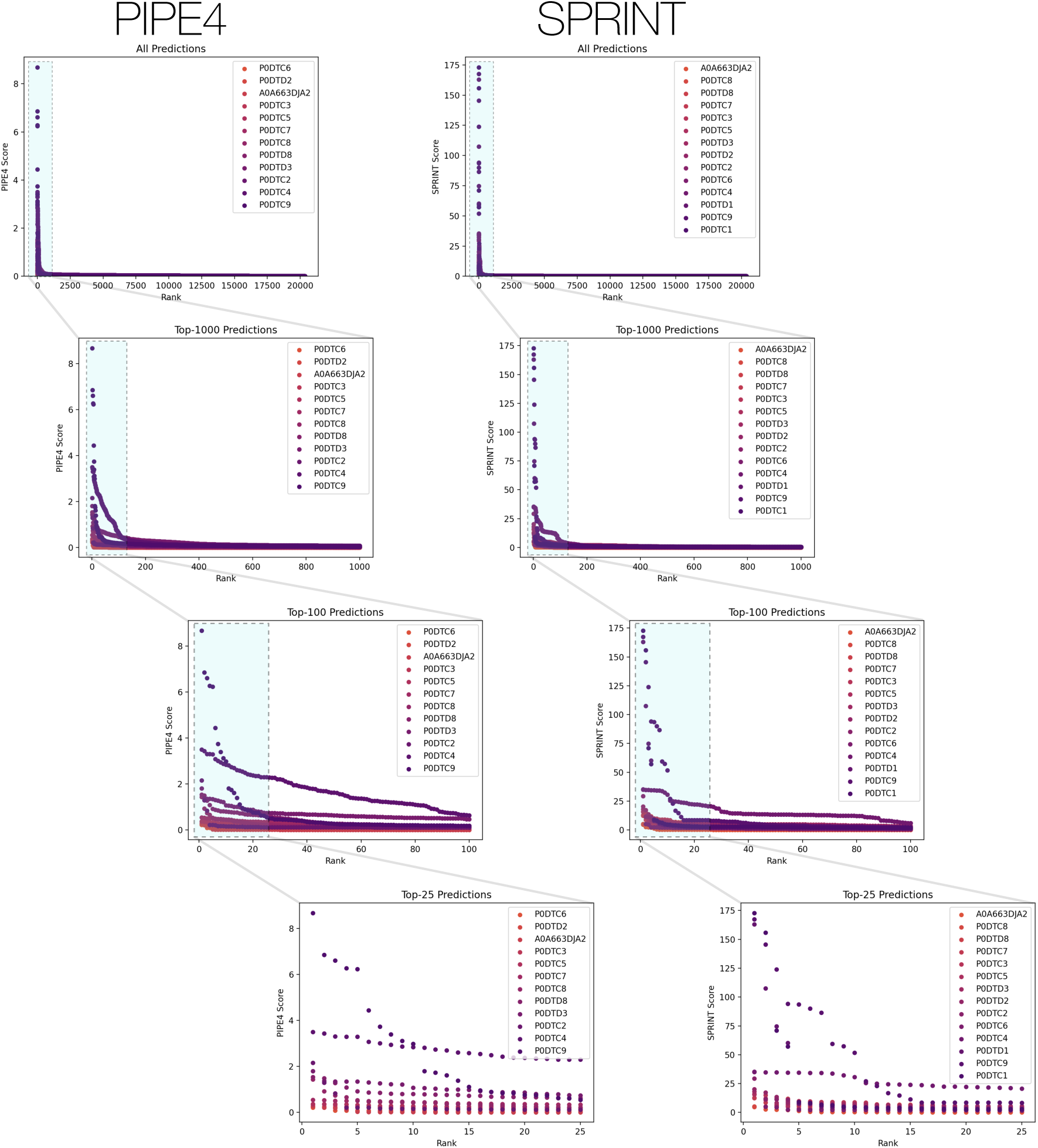
One-to-All Score Curves by Top-*k*. The top panels depict the combination of one-to-all score curves for each protein, by each predictor and each subplot is a top-*k* subset of the previous; highlighted in blue.

We automated the selection of this operational decision threshold for the 14 SARS-CoV-2 proteins using the Kneedle algorithm, applied to its top-1000 predictions, using a sensitivity parameter of 2.0. An example visual illustration of the highly conservative selection of high-confidence interactions is depicted in Figure 4 and the cut-offs for each protein are tabulated in Table 3.

**Figure 3:**
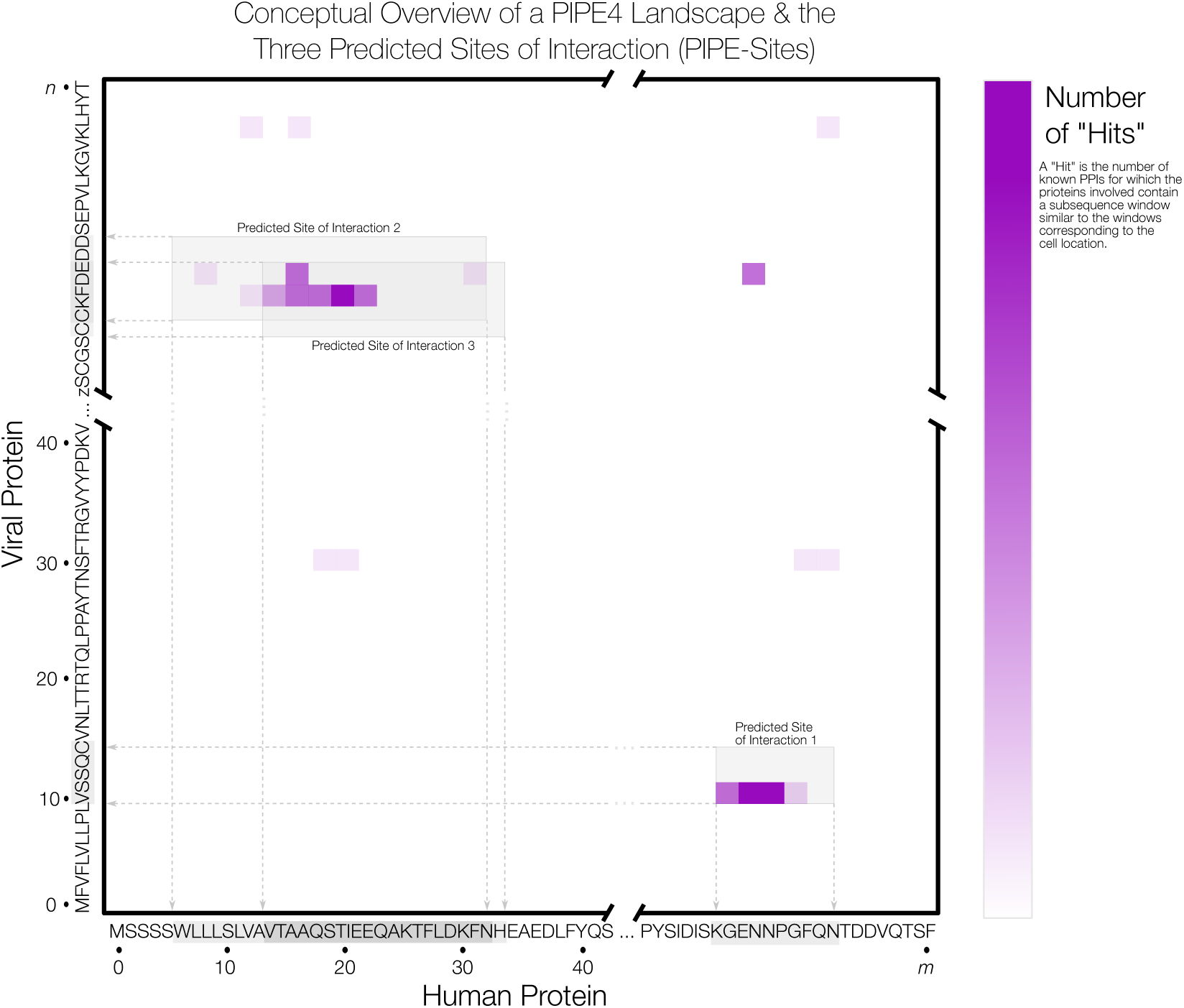
Conceptual Overview of the PIPE4 Landscape and the Three Predicted Sites of Interaction (PIPE-Sites)

**Figure 4:**
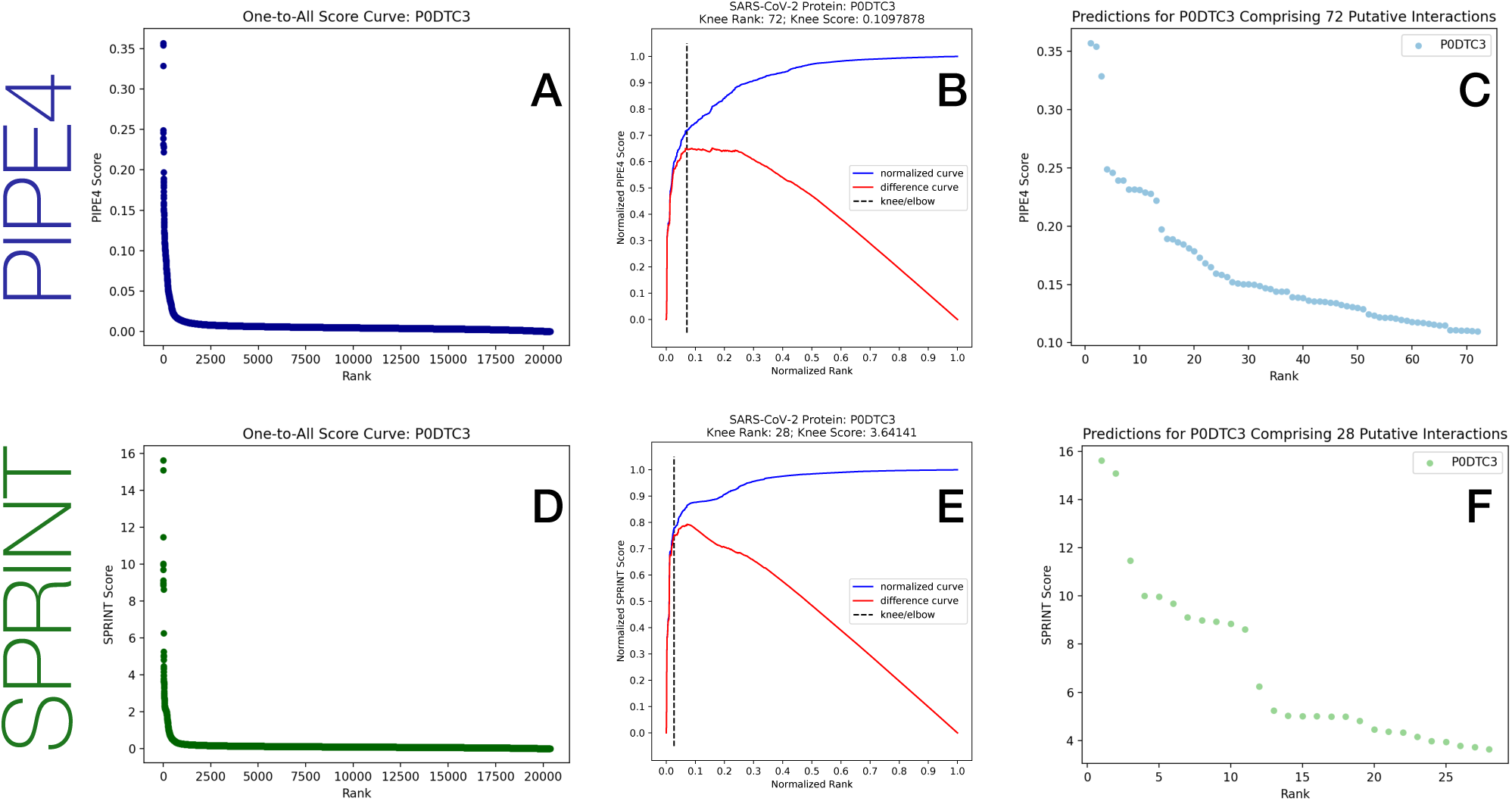
Example Compilation of the Spike Protien One-to-All Score Curve, Knee Detection for Local Cut-Off, and Rank Order Predictions, for each Method. Panels A & D depict the one-to-all score curves from the predicted score between the Spike protein and all proteins in the human proteome. Panels B & E depict the detected knee from the top-1000 of the one-to-all score curves. Panels C & F depict the predicted interactions above the knee.

We identified the common set of predicted pairs above each locally defined knee from *both* the PIPE4 and SPRINT methods (their intersection), resulting in a set of 225 putative human protein targets among 279 intersection pairs. These pairs were considered the predicted interactome and were subsequently analyzed by PIPE-Sites and GO-term enrichment analysis was performed using the 225 human proteins identifed.

#### 2.2.4 Predicted PPI Site of Interaction using PIPE-Sites

The PIPE4 algorithm generates its prediction for a given pair of proteins based on a two-dimensional landscape of scores, where the score at location *x, y*, the number of sequence window similarity “hits”, represents the weight of evidence from the *x*^*th*^ and *y*^*th*^ subsequence of the human and SARS-CoV-2 proteins, respectively. The PIPE-Sites algorithm examines this landscape and deduces which subsequences from each protein are likely to correspond to the site of interaction [8]. Such information can guide subsequent detailed investigations to determine the physical binding site which may form the target for novel interventions to disrupt the PPI.

The list of PPIs generated from both methods can be used to inform the design of anti-SARS-CoV-2 therapeutics by using peptide sequences from the predicted PPI site, which we refer to as the PPI-Site. We define the PPI-Site as the peptide sequence that is responsible for mediating a given PPI, which is here estimated using the PIPE-Sites method. A conceptual overview of the PIPE4 landscape matrix and PIPE-Site prediction is illustrated in Figure 3.

### 2.3 Gene Ontology (GO) Enrichment Analysis

To determine which cellular pathways may be targeted by SARS-CoV-2, PANTHER Gene Ontology (GO) enrichment analysis was applied to all 225 human proteins predicted to interact with SARS-COV-2 proteins. The molecular function, biological pathway, and cellular pathway *p*-values were determined with the Fisher’s Exact test implemented in the PANTHER Go software [12]. P-Values were corrected for multiple testing using the False Discovery Rate (FDR) method [17] and significant terms were identified at a threshold of 0.05.

### 2.4 High-Performance Computing Infrastructure

In order to generate the ∼280,000 PPI predictions, high-performance computing infrastructure was required. The ComputeCanada *Graham* heterogeneous cluster was leveraged to generate these predictions. Boasting more than 41,000 cores and 520 GPU devices across 1,185 nodes, this HPC cluster enabled the rapid computation and compilation of these predictions. Computational research related to the COVID-19 pandemic has been assigned increased priority which expedited the generation of these predictions.

## 3 Results & Discussion

It is of critical importance that the global research community focus its efforts on the rapid understanding the SARS-CoV-2 virus and the pathogenesis of COVID-19 in order to develop anti-viral therapeutics. Fortunately, the prior decades of research into related viral families provide a wealth of data with which to guide current and future studies, such as with the elucidation of the SARS-CoV vs human inter-species interactome in 2011 using the high-throughput (though false positive-prone) yeast-two hybrid method to highlight cyclophilins as a target for pan-coronavirus inhibitors [18]. Previous knowledge of related coronaviruses within the *Coronaviridae* family provide training samples with which we can identify a number of new high-confidence PPIs that contribute to our understanding of the COVID-19 disease pathogenesis and which may represent targets for novel inhibitory therapeutics.

Notably, it is known that the SARS-CoV Spike protein binds to the human ACE2 receptor [19]. Upon entry into the respiratory or gastrointestinal tracts, coronaviruses establish themselves by entering and infecting lumenal macrophages and epithelial cells. The viral cell entry program is orchestrated by the spike protein that binds to the human cellular receptors and, thereby, mediates virus-cell membrane fusions.

While the putative interaction between the SARS-CoV-2 Spike protein and human ACE2 receptor is a current focus of the research community, it is also valuable to develop a more holistic understanding of the possibly numerous SARS-CoV-2 vs. human PPIs. Consequently, additional viral-human interactions might be targeted and disrupted with the use of small inhibitory peptides or molecules. To this end, we leverage sequence-based predictors to score all possible interactions between the SARS-CoV-2 and human proteomes. For each of the 14 viral proteins, sorting the 20,366 scores (for each human protein) into a monotonically decreasing rank order enables the identification of the subset of high scoring putative interactors with that one viral protein (Figure 4A,D).

Rather than apply a globally defined decision threshold (*i*.*e*. top-*k* or minimum threshold), we automatically detected a highly conservative “knee” for each curve (the point of greatest rate of change) to delineate those rare high-scoring pairs from the remaining baseline (Figure 4B,E). The union of the *n* = 1210 predicted PIPE4 and SPRINT high confidence putative PPIs comprises only ∼0.42% of all possible pairs, and their intersection of *n* = 279 putative pairs comprises a highly conservative <0.098%. These data are tabulated in Table3, illustrated in Figure 5, and plotted in Figure 4.

**Figure 5:**
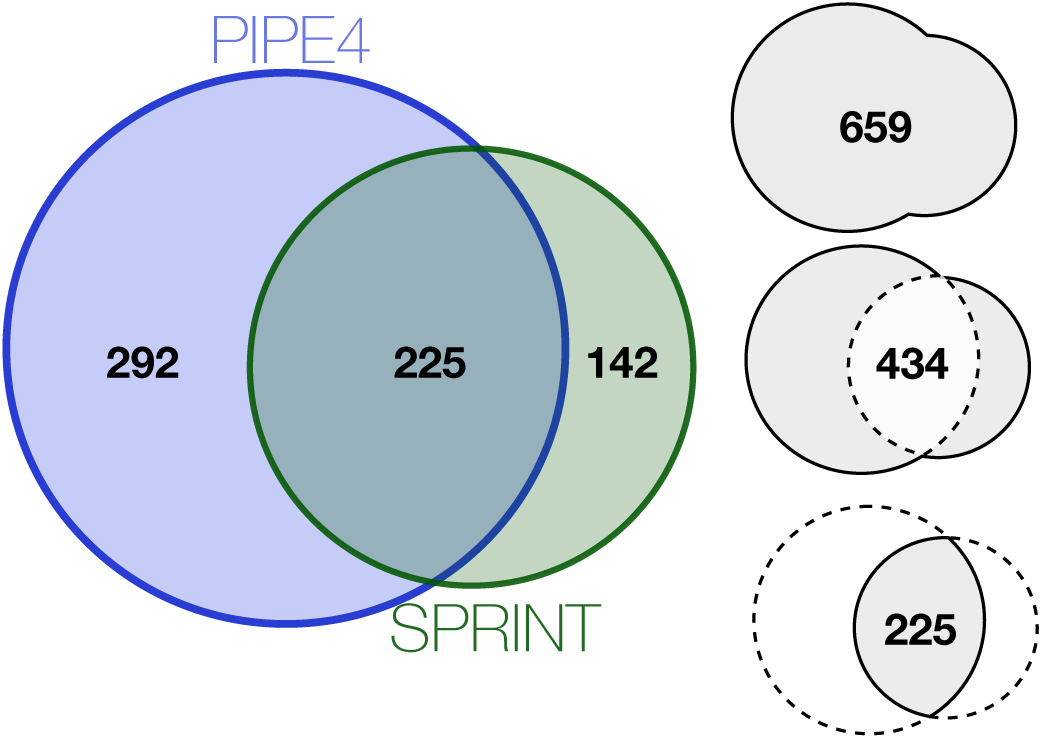
Venn Diagram of the Human Proteins Predicted to Interact with SARS-CoV-2 Proteins.

We provide the landscapes and predicted PIPE-Sites for each of the predicted interactions for each SARS-CoV-2 proteins and highlight those 279 pairs within the predicted interactome. All data are published in the following DataVerse repository, [10].

### 3.1 On the Interpretation of PIPE-Sites Predictions

When interpreting the landscapes, it is important to note that the PIPE-Sites algorithm used here is simplistic in its implementation. Briefly, a maximum of three potential peaks in the landscape are identified and a walk algorithm expands the predicted site of interaction until the score falls below a given threshold [8].

The highlighted sites may appear “shifted” relative to the highlighted cells (typically in the bottom-left); this is due to the algorithm’s use of a window of 20 amino acids in length that extends both to the left (along the x-axis) and upwards (along the y-axis). Consequently, the *minimum PIPE-Site size* is 20 × 20 with the peak in the bottom-left corner. Additionally, this implementation may result in the predicted site extending past the coloured matrix, either to the right or above. This defined window size additionally prevents predictions within the terminal 20 amino acids of both sequences given that the widow sizes in these regions would necessarily be less than 20 amino acids in length. Finally, the PIPE-Sites may overlap when numerous hits appear within close proximity, as is the case when a “band” of hits appears in the matrix. Finally, when the peak of the landscape comprises only a few hits (generally <3) the entire landscape is predicted as a site of interaction; evidently, these should be disregarded.

Therefore, when interpreting the landscapes, it is important not to solely rely on these proposed regions; they function as an initial guide, yet other high-scoring, or “hot-spot”, regions of interest may exist in the landscape. By providing a matrix of raw scores (in the form of a space-separated .mat file), visual interpretation of the results promise to reveal notable subsequences as well as enable the application of related interaction site predictors to identify putative sites of interaction.

### 3.2 The Spike Protein vs. ACE2 Interaction

Most excitingly, both the PIPE4 and SPRINT predictors scored the SARS-CoV-2 Spike protein vs. human ACE2 protein as the top-ranking prediction in their respective one-to-all score curves (P0DTC2-Q9BYF1) (PIPE4 SW score of 2.159, SPRINT score of 29.3515). As previously noted, this was achieved despite the removal of the known SARS-CoV Spike protein vs. ACE2 PPI within the training dataset. This finding corroborates related research reporting that SARS-CoV-2 can infect the human respiratory epithelial cells through interaction with the human ACE2 receptor [20].

Certainly, if the SARS-CoV-2 and SARS-CoV Spike proteins share sufficient sequence and structural similairty, it can be expected that anti-virals designed against SARS-CoV promise to also be effective against SARS-CoV-2. We investigate this sequence similarity by performing a BLASTp alignment of the two sequences. Interestingly, only 76% identity was observed (Figure 7) suggesting that the SARS-CoV-2 protein might have evolved to be sufficiently different from its SARS-CoV variant to render existing anti-virals ineffective. However, the SARS-CoV-2 variant is likely to share a similar mechanism of action where the recombinant SARS-CoV-2 spike protein downregulates ACE2 expression and thereby promotes lung injury, as seen in the SARS-CoV variant [19].

**Figure 6:**
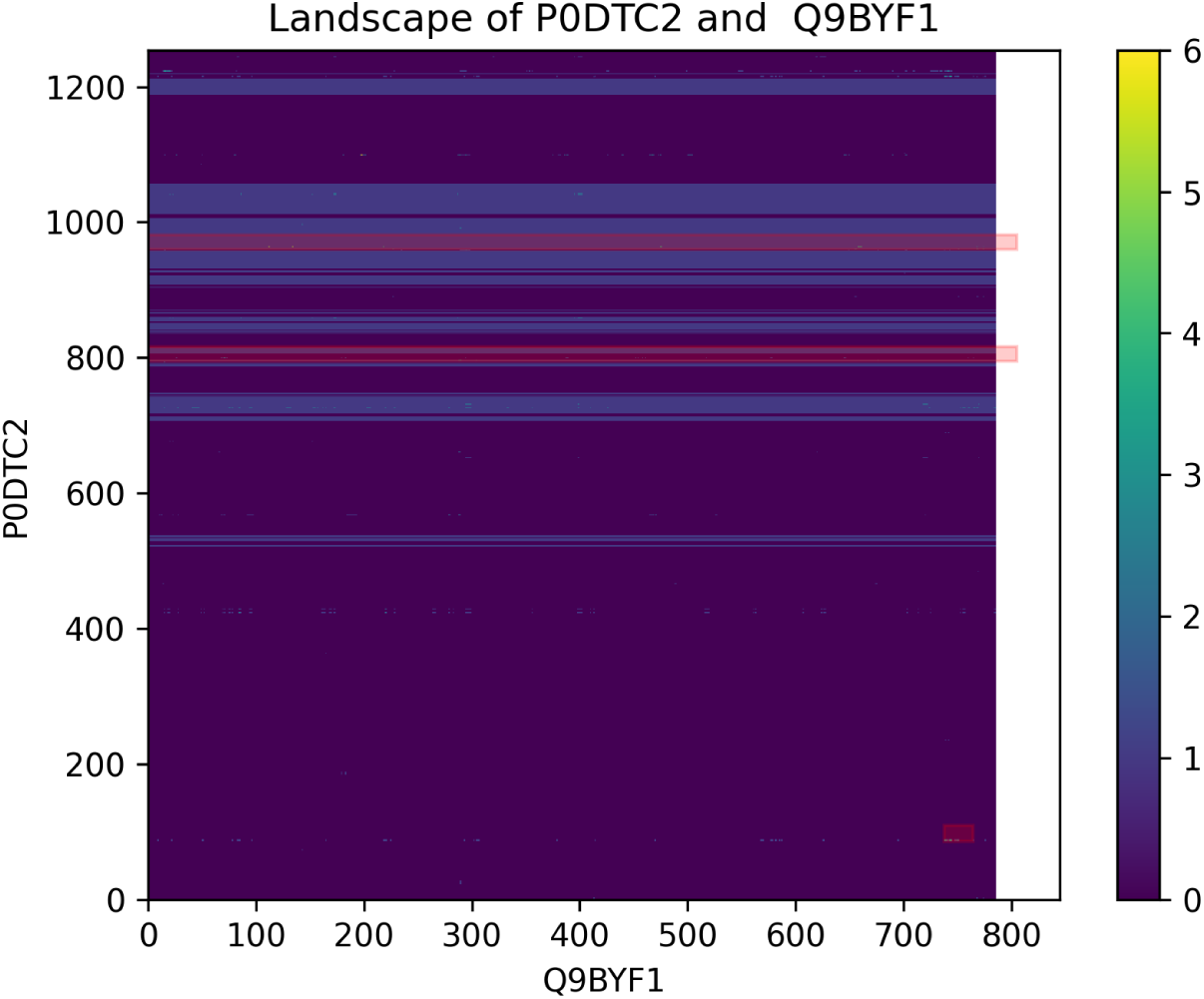
The PIPE-Sites Landscape between the SARS-CoV-2 Spike protein and human ACE2 protein. The three red rectangles represent the predicted PIPE-Sites regions.

**Figure 7:**
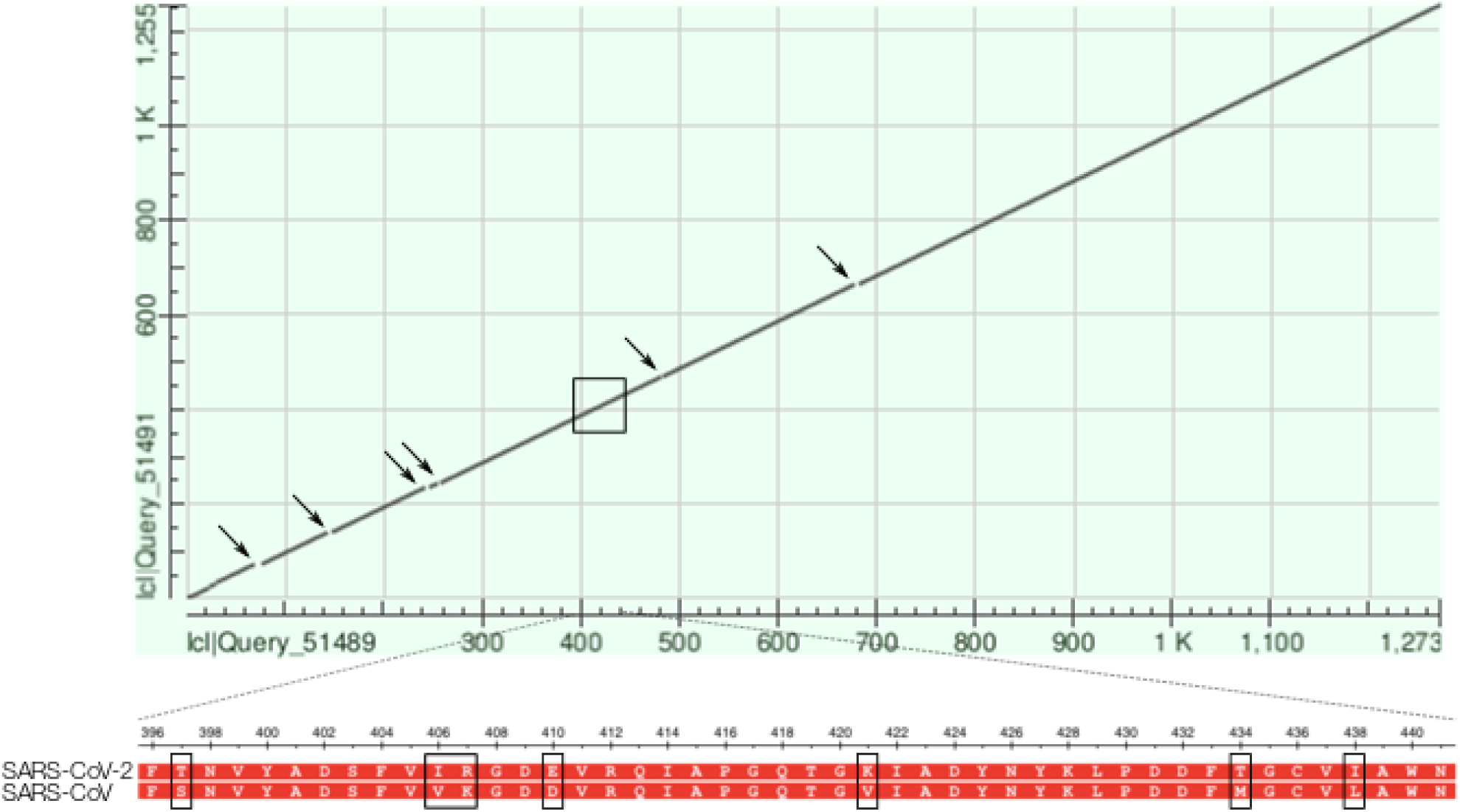
Dot Plot of the BLASTp Alignment of the SARS-CoV and SARS-CoV-2 Spike Protein. The alignment of the two proteins results in a *max score* of 2039, a *total score* of 2039, 100% coverage, an *E-value* of 0.0, and 76.04% identity. Specifically: 971/1277 (76%) identities, 1109/1277 (86%) positives, and 26/1277 (2%) gaps. Arrows indicate gaps within the alignment and the zoomed-in region highlights the six mismatches around residue 420.

Consequently, the elucidation of the Spike-ACE2 binding interface is needed to design novel therapeutics. To that end, we used the PIPE-Sites algorithm to predict the three most likely putative interaction interfaces between the Spike (P0DTC2) and ACE2 (Q9BYF1) proteins (Figure 6). Note that all predicted subsequence offsets are 0-indexed. With a maximum landscape peak of 6, the PIPE-Sites algorithm identified three putative interaction interfaces:

1. **P0DTC2**: [86–109]; **Q9BYF1**: [738–816]
2. **P0DTC2**: [795–816]; **Q9BYF1**: Entire sequence
3. **P0DTC2**: [960–981]; **Q9BYF1**: Entire sequence

Interestingly, the PIPE-Sites score landscape in Figure 6 exhibits a number of horizontal bands indicative of subsequence regions along the Spike protein that correspond to a relatively high likelihood of interaction. While the PIPE-Sites algorithm only identifies three putative regions, these bands suggest additional regions of interest.

The highest-scoring predicted PIPE-Site interface corresponds to the Spike [86–109] subsequence and the ACE2 [738–816] subsequence, which resides within the *intracellular* cytoplasmic domain of ACE2. However, upon closer inspection of other “hot spot” regions within the landscape, we note several that reside within the *extracellular* N-terminal region of ACE2 (*i*.*e*. residues ∼[30-84] & [353-357]). In particular, we note the three following regions of interest:

> *Visually high-scoring region:* **P0DTC2**: near residue 1224; **Q9BYF1**: [15–23]
>
> *Within ACE2 residues [30-84]:* **P0DTC2**: near residue 420; **Q9BYF1**: [80-84]
>
> *Within ACE2 residues [353-537]:* **P0DTC2**: near residue 420; **Q9BYF1**: [355-357]

Most interestingly, certain of these region along the Spike protein appears to coincide with mismatched or gap regions along the dot plot alignment depicted in Figure 7). For example, upon closer investigation of the alignment around residue 420, we note six mismatches. Their proximity to a candidate region of interaction certainly warrant additional experimental investigation (Figure 7).

While numerous inhibitory strategies exist, including the use of small molecules or small interfering RNAs, this research is most directly amenable to the design small inhibitory peptides that inhibit virus infection by preventing Spike protein-mediated receptor binding and blocking viral fusion and entry. Unfortunately, much like small peptides and interfering RNAs, peptide-based solutions are disadvantaged by their *low antiviral potency*.

### 3.3 HLA Class I/II Histocompatibility Antigen

Among the 225 human proteins identified, six Human Leukocyte Antigen (HLA) class I/II histocompatibility antigens were predicted to interact with P0DTC3, the SARS-CoV-2 Protein 3a:

- **P13747**: HLA-E HLA-6.2 HLAE
- **P01911**: HLA-DRB1
- **P17693**: HLA-G HLA-6.0 HLAG
- **P04439**: HLA-A HLAA
- **P10321**: HLA-C HLAC
- **P30511**: HLA-F HLA-5.4 HLAF

The visualization of the predicted site of interaction for the six HLA interactions highlight a consistent subsequence region of the SARS-CoV-2 protein 3a between amino acids [202–222] (Figure 8). Literature review reveals that the open reading frames (ORFs) of the SARS-CoV virus, the ORF3a, encodes the variant 274 AA-long Protein 3a. A previous study used sequence analysis that suggested that the ORF3a aligned to a calcium pump present in *Plasmodium falciparum* and glutamine synthetase found in *Leptospira interrogans*. This sequence similarity between the three organisms was found to be limited only to amino acid residues [209–264] which form the cytoplasmic domain of ORF3a. This subsequence region was predicted to be involved in calcium binding and then confirmed *in vitro* [21].

**Figure 8:**
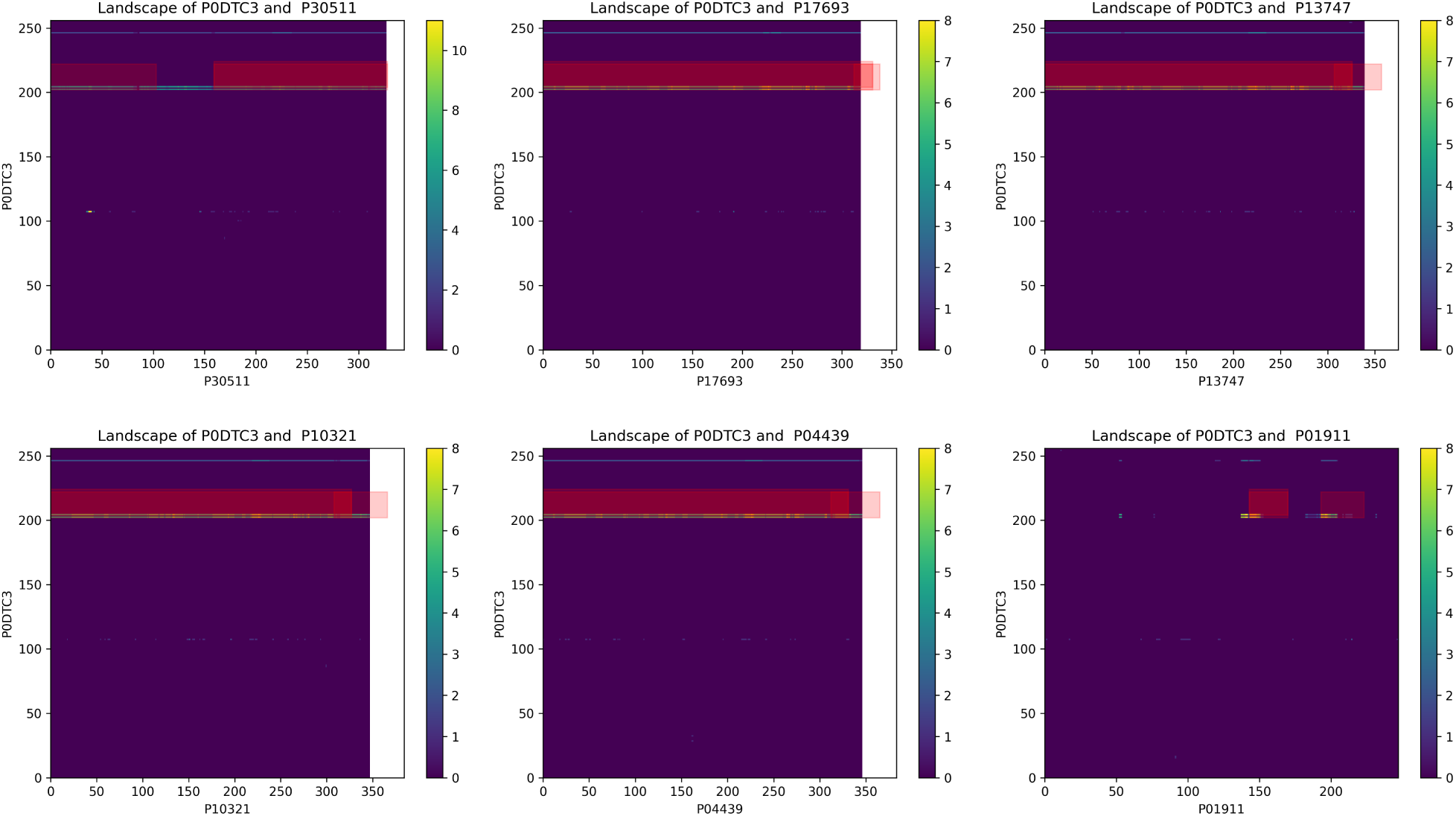
Landscapes of the Six Predicted HLA interactors with SARS-CoV-2 Protein 3a. The three red rectangles represent the predicted PIPE-Sites regions. Their “shifted” relative to the highlighted cells is due to the algorithm’s use of a window of 20 amino acids in length that extends both to the left (along the x-axis) and upwards (along the y-axis). This implementation may also result in the predicted site extending past the coloured matrix, either to the right or above. The PIPE-Sites may overlap when numerous hits appear within close proximity, as is the case when a “band” of hits appears in the matrix.

Given the important role that calcium plays as part of virion structure formation, virus entry, viral gene expression, virion maturation, and release, these regions of Protein 3a are of possible interest for disruption of SARS-CoV-2. Specifically, the design of a small inhibitory peptide targeting this subsequence region of protein 3a might disrupt the viral life cycle.

### 3.4 GO-Term Analyses

Of the 225 human proteins within the intersection of the PIPE4 and SPRINT predicted interactions, we ran a number of GO-term analyses to better understand the functional role of the human proteins involved. To this end, the GO Panther Classification System was used to run over/under-representation analysis of the 225 human proteins as compared to the reference human proteome. A Fisher’s Exact test with correction for False Discovery Rate was used to extract a list of the most enriched GO-terms among the 215 human proteins for which GO-term data were available. The Molecular Functions exhibiting a fold enrichment greater than 3 are reported in 5; the Biological Processes exhibiting a fold enrichment greater than 50 are reported in 6; and the Cellular Components exhibiting a fold enrichment greater than 15 are reported in 7. The fold enrichment cut-offs were selected to limit the size of the tables; the complete tables are available in the public repository, [10].

While this current analysis combines all predicted human interactors together, a more revealing analysis might investigate the resultant GO-terms on a per-viral-protein basis to identify those human pathways and biological processes most sensibly targeted by SARS-CoV-2. This analysis is likely to appear in a subsequent version of this work.

Of the GO-terms observed from the current analysis, the highly over-represented biological processes in Table 6 are the most interesting. Notably, the top-9 GO-terms have a 96.98 fold enrichment given that the predicted set of human interactors contain all of the proteins from the *H. sapiens* reference (*i*.*e*. the number of proteins present in the reference are also in the sample: 2/2, 8/8, and 3/3 among the top-3, respectively). We specifically highlight the “antigen processing and presentation of exogenous peptide antigen via MHC class Ib” (GO:0002477) and the “calcium ion transport from cytosol to endoplasmic reticulum” (GO:1903515). Moreover, the top-ranking cellular component GO-terms (Table 7) show notable over-representation of “MHC class Ib protein complex” (GO:0032398), “MHC class I protein complex” (GO:0042612), and numerous proteasome complex terms. While only a shallow analysis is presented here, a more involved investigation into these predicted interactions promises to reveal putative targets for novel inhibitory peptides.

**Table 5:** PANTHER GO-Term Analysis of Molecular Function Over/Under-Representation for the 225 Predicted Human Interactors.

| PANTHER GO Molecular Function | Homo sapiens<br>Reference (N=20,851) | Num.<br>Predicted | Predicted<br>Num. Expected | Over/Under<br>Represented | Fold<br>Enrichment | p-value | FDR |
| --- | --- | --- | --- | --- | --- | --- | --- |
| peptidase activator activity (GO:0016504) | 6 | 4 | 0.06 | + | 64.65 | 2.11E-06 | 4.88E-05 |
| tumor necrosis factor receptor superfamily binding<br>(GO:0032813) | 9 | 5 | 0.09 | + | 53.88 | 1.96E-07 | 6.53E-06 |
| TBP-class protein binding (GO:0017025) | 9 | 3 | 0.09 | + | 32.33 | 2.15E-04 | 3.28E-03 |
| ubiquitin-like protein ligase binding (GO:0044389) | 46 | 14 | 0.47 | + | 29.52 | 9.96E-16 | 1.77E-13 |
| protein tyrosine kinase activity (GO:0004713) | 61 | 17 | 0.63 | + | 27.03 | 2.71E-18 | 7.20E-16 |
| ubiquitin protein ligase binding (GO:0031625) | 41 | 9 | 0.42 | + | 21.29 | 1.77E-09 | 1.05E-07 |
| signal sequence binding (GO:0005048) | 33 | 7 | 0.34 | + | 20.57 | 1.47E-07 | 6.00E-06 |
| heat shock protein binding (GO:0031072) | 30 | 5 | 0.31 | + | 16.16 | 2.67E-05 | 4.91E-04 |
| ATP binding (GO:0005524) | 40 | 5 | 0.41 | + | 12.12 | 9.26E-05 | 1.54E-03 |
| unfolded protein binding (GO:0051082) | 58 | 6 | 0.6 | + | 10.03 | 4.83E-05 | 8.28E-04 |
| endopeptidase activity (GO:0004175) | 307 | 24 | 3.17 | + | 7.58 | 6.01E-14 | 5.33E-12 |
| ATPase activity, coupled (GO:0042623) | 117 | 8 | 1.21 | + | 6.63 | 4.37E-05 | 7.75E-04 |
| peptidase activity (GO:0008233) | 415 | 28 | 4.28 | + | 6.54 | 1.35E-14 | 1.43E-12 |
| peptidase activity, acting on L-amino acid peptides<br>(GO:0070011) | 407 | 27 | 4.2 | + | 6.43 | 6.06E-14 | 4.61E-12 |
| ubiquitin-protein transferase activity (GO:0004842) | 239 | 14 | 2.46 | + | 5.68 | 3.57E-07 | 1.00E-05 |
| peptide binding (GO:0042277) | 194 | 11 | 2 | + | 5.5 | 8.82E-06 | 1.80E-04 |
| ubiquitin-like protein transferase activity<br>(GO:0019787) | 249 | 14 | 2.57 | + | 5.45 | 5.70E-07 | 1.52E-05 |
| ubiquitin protein ligase activity (GO:0061630) | 145 | 8 | 1.5 | + | 5.35 | 1.80E-04 | 2.91E-03 |
| cytokine receptor binding (GO:0005126) | 93 | 5 | 0.96 | + | 5.21 | 3.33E-03 | 4.43E-02 |
| ubiquitin-like protein ligase activity (GO:0061659) | 149 | 8 | 1.54 | + | 5.21 | 2.15E-04 | 3.36E-03 |
| amide binding (GO:0033218) | 211 | 11 | 2.18 | + | 5.06 | 1.86E-05 | 3.67E-04 |
| catalytic activity, acting on a protein (GO:0140096) | 1400 | 65 | 14.44 | + | 4.5 | 5.29E-25 | 2.81E-22 |
| ATPase activity (GO:0016887) | 262 | 12 | 2.7 | + | 4.44 | 2.65E-05 | 5.04E-04 |
| phosphotransferase activity, alcohol group as acceptor<br>(GO:0016773) | 519 | 21 | 5.35 | + | 3.92 | 1.77E-07 | 6.28E-06 |
| protein kinase activity (GO:0004672) | 435 | 17 | 4.49 | + | 3.79 | 4.31E-06 | 9.56E-05 |
| enzyme binding (GO:0019899) | 610 | 23 | 6.29 | + | 3.66 | 1.50E-07 | 5.71E-06 |
| kinase activity (GO:0016301) | 559 | 21 | 5.76 | + | 3.64 | 5.71E-07 | 1.45E-05 |
| signaling receptor binding (GO:0005102) | 629 | 23 | 6.49 | + | 3.55 | 2.53E-07 | 7.49E-06 |
| transferase activity, transferring phosphorus-containing<br>groups (GO:0016772) | 665 | 21 | 6.86 | + | 3.06 | 7.94E-06 | 1.69E-04 |

**Table 6:** PANTHER GO-Term Analysis of Biological Process Over/Under-Representation for the 225 Predicted Human Interactors.

| PANTHER GO Biological Process | Homo sapiens<br>Reference (N=20,851) | Num.<br>Predicted | Predicted<br>Num. Expected | Over/Under<br>Represented | Fold<br>Enrichment | p-value | FDR |
| --- | --- | --- | --- | --- | --- | --- | --- |
| antigen processing and presentation of exogenous peptide antigen via MHC class Ib (GO:0002477) | 2 | 2 | 0.02 | + | 96.98 | 6.14E-04 | 1.53E-02 |
| nerve growth factor production (GO:0032902) | 2 | 2 | 0.02 | + | 96.98 | 6.14E-04 | 1.53E-02 |
| neurotrophin production (GO:0032898) | 2 | 2 | 0.02 | + | 96.98 | 6.14E-04 | 1.52E-02 |
| positive regulation of endoplasmic reticulum calcium ion concentration (GO:0032470) | 2 | 2 | 0.02 | + | 96.98 | 6.14E-04 | 1.52E-02 |
| entry of viral genome into host nucleus through nuclear pore complex via importin (GO:0075506) | 2 | 2 | 0.02 | + | 96.98 | 6.14E-04 | 1.52E-02 |
| positive regulation of telomerase RNA reverse transcriptase activity (GO:1905663) | 2 | 2 | 0.02 | + | 96.98 | 6.14E-04 | 1.52E-02 |
| positive regulation of fast-twitch skeletal muscle fiber contraction (GO:0031448) | 2 | 2 | 0.02 | + | 96.98 | 6.14E-04 | 1.51E-02 |
| regulation of fast-twitch skeletal muscle fiber contraction (GO:0031446) | 2 | 2 | 0.02 | + | 96.98 | 6.14E-04 | 1.51E-02 |
| calcium ion transport from cytosol to endoplasmic reticulum (GO:1903515) | 2 | 2 | 0.02 | + | 96.98 | 6.14E-04 | 1.51E-02 |
| multi-organism nuclear import (GO:1902594) | 4 | 3 | 0.04 | + | 72.74 | 3.56E-05 | 1.18E-03 |
| viral penetration into host nucleus (GO:0075732) | 4 | 3 | 0.04 | + | 72.74 | 3.56E-05 | 1.18E-03 |
| nerve growth factor processing (GO:0032455) | 4 | 3 | 0.04 | + | 72.74 | 3.56E-05 | 1.18E-03 |
| adenine transport (GO:0015853) | 4 | 3 | 0.04 | + | 72.74 | 3.56E-05 | 1.18E-03 |
| proteasomal ubiquitin-independent protein catabolic process (GO:0010499) | 23 | 16 | 0.24 | + | 67.47 | 2.42E-22 | 3.01E-20 |
| histamine secretion by mast cell (GO:0002553) | 3 | 2 | 0.03 | + | 64.65 | 1.02E-03 | 2.33E-02 |
| histamine secretion involved in inflammatory response (GO:0002441) | 3 | 2 | 0.03 | + | 64.65 | 1.02E-03 | 2.33E-02 |
| positive regulation of caveolin-mediated endocytosis (GO:2001288) | 3 | 2 | 0.03 | + | 64.65 | 1.02E-03 | 2.33E-02 |
| histamine production involved in inflammatory response (GO:0002349) | 3 | 2 | 0.03 | + | 64.65 | 1.02E-03 | 2.32E-02 |
| regulation of telomerase RNA reverse transcriptase activity (GO:1905661) | 3 | 2 | 0.03 | + | 64.65 | 1.02E-03 | 2.32E-02 |
| positive regulation of translation in response to endoplasmic reticulum stress (GO:0036493) | 3 | 2 | 0.03 | + | 64.65 | 1.02E-03 | 2.32E-02 |
| calcium ion import into sarcoplasmic reticulum (GO:1990036) | 3 | 2 | 0.03 | + | 64.65 | 1.02E-03 | 2.31E-02 |
| positive regulation of ATPase-coupled calcium transmembrane transporter activity (GO:1901896) | 5 | 3 | 0.05 | + | 58.19 | 5.65E-05 | 1.80E-03 |

**Table 7:** PANTHER GO-Term Analysis of Cellular Component Over/Under-Representation for the 225 Predicted Human Interactors.

| PANTHER GO Cellular Component | Homo sapiens<br>Reference (N=20,851) | Num.<br>Predicted | Predicted<br>Num. Expected | Over/Under<br>Represented | Fold<br>Enrichment | p-value | FDR |
| --- | --- | --- | --- | --- | --- | --- | --- |
| MHC class Ib protein complex (GO:0032398) | 2 | 2 | 0.02 | + | 96.98 | 6.14E-04 | 1.13E-02 |
| proteasome core complex, alpha-subunit complex (GO:0019773) | 8 | 8 | 0.08 | + | 96.98 | 1.24E-12 | 8.33E-11 |
| proteasome activator complex (GO:0008537) | 3 | 3 | 0.03 | + | 96.98 | 2.05E-05 | 5.04E-04 |
| spermatoproteasome complex (GO:1990111) | 5 | 4 | 0.05 | + | 77.59 | 1.28E-06 | 4.02E-05 |
| phosphopyruvate hydratase complex (GO:0000015) | 4 | 3 | 0.04 | + | 72.74 | 3.56E-05 | 8.64E-04 |
| proteasome core complex (GO:0005839) | 21 | 15 | 0.22 | + | 69.27 | 3.82E-21 | 5.50E-19 |
| proteasome core complex, beta-subunit complex (GO:0019774) | 11 | 7 | 0.11 | + | 61.72 | 3.03E-10 | 1.60E-08 |
| proteasome regulatory particle, base subcomplex (GO:0008540) | 12 | 7 | 0.12 | + | 56.57 | 4.75E-10 | 2.33E-08 |
| MHC class I protein complex (GO:0042612) | 9 | 5 | 0.09 | + | 53.88 | 1.96E-07 | 6.82E-06 |
| eukaryotic translation elongation factor 1 complex (GO:0005853) | 4 | 2 | 0.04 | + | 48.49 | 1.51E-03 | 2.44E-02 |
| cytosolic proteasome complex (GO:0031597) | 9 | 4 | 0.09 | + | 43.1 | 7.02E-06 | 1.94E-04 |
| protein phosphatase type 1 complex (GO:0000164) | 9 | 4 | 0.09 | + | 43.1 | 7.02E-06 | 1.91E-04 |
| PTW/PP1 phosphatase complex (GO:0072357) | 7 | 3 | 0.07 | + | 41.56 | 1.19E-04 | 2.67E-03 |
| proteasome complex (GO:0000502) | 65 | 26 | 0.67 | + | 38.79 | 8.25E-31 | 3.32E-28 |
| proteasome accessory complex (GO:0022624) | 25 | 10 | 0.26 | + | 38.79 | 1.46E-12 | 9.49E-11 |
| endopeptidase complex (GO:1905369) | 66 | 26 | 0.68 | + | 38.2 | 1.14E-30 | 3.83E-28 |
| CD40 receptor complex (GO:0035631) | 11 | 4 | 0.11 | + | 35.27 | 1.32E-05 | 3.36E-04 |
| platelet dense tubular network membrane (GO:0031095) | 9 | 3 | 0.09 | + | 32.33 | 2.15E-04 | 4.62E-03 |
| cytoplasmic side of lysosomal membrane (GO:0098574) | 6 | 2 | 0.06 | + | 32.33 | 2.79E-03 | 4.01E-02 |
| proteasome regulatory particle (GO:0005838) | 22 | 7 | 0.23 | + | 30.86 | 1.35E-08 | 5.91E-07 |
| peptidase complex (GO:1905368) | 91 | 26 | 0.94 | + | 27.71 | 1.18E-27 | 2.64E-25 |
| glycogen granule (GO:0042587) | 7 | 2 | 0.07 | + | 27.71 | 3.56E-03 | 4.98E-02 |
| platelet dense tubular network (GO:0031094) | 11 | 3 | 0.11 | + | 26.45 | 3.51E-04 | 7.01E-03 |
| pseudopodium (GO:0031143) | 17 | 4 | 0.18 | + | 22.82 | 5.51E-05 | 1.31E-03 |
| postsynaptic specialization, intracellular component (GO:0099091) | 22 | 5 | 0.23 | + | 22.04 | 7.11E-06 | 1.91E-04 |
| integral component of luminal side of endoplasmic reticulum membrane (GO:0071556) | 29 | 6 | 0.3 | + | 20.07 | 1.34E-06 | 4.14E-05 |
| luminal side of endoplasmic reticulum membrane (GO:0098553) | 29 | 6 | 0.3 | + | 20.07 | 1.34E-06 | 4.08E-05 |
| MHC protein complex (GO:0042611) | 28 | 5 | 0.29 | + | 17.32 | 1.99E-05 | 4.94E-04 |
| COP9 signalosome (GO:0008180) | 36 | 6 | 0.37 | + | 16.16 | 4.07E-06 | 1.21E-04 |
| luminal side of membrane (GO:0098576) | 36 | 6 | 0.37 | + | 16.16 | 4.07E-06 | 1.19E-04 |
| extrinsic component of cytoplasmic side of plasma membrane (GO:0031234) | 93 | 15 | 0.96 | + | 15.64 | 3.15E-13 | 2.19E-11 |

### 3.5 Complete Predicted Interactome

To better visualize the predicted interactome and the over-represented GO-terms within, a network-based representation is depicted in Figure 9. Much like the HLA proteins highlighted above, we note a number of highly represented GO-terms around several of the proteins of interest including those related to the immune response, various types of signalling, and the viral life cycle. We hope that this work will guide the broader research community in their search for putative inhibitory molecules.

**Figure 9:**
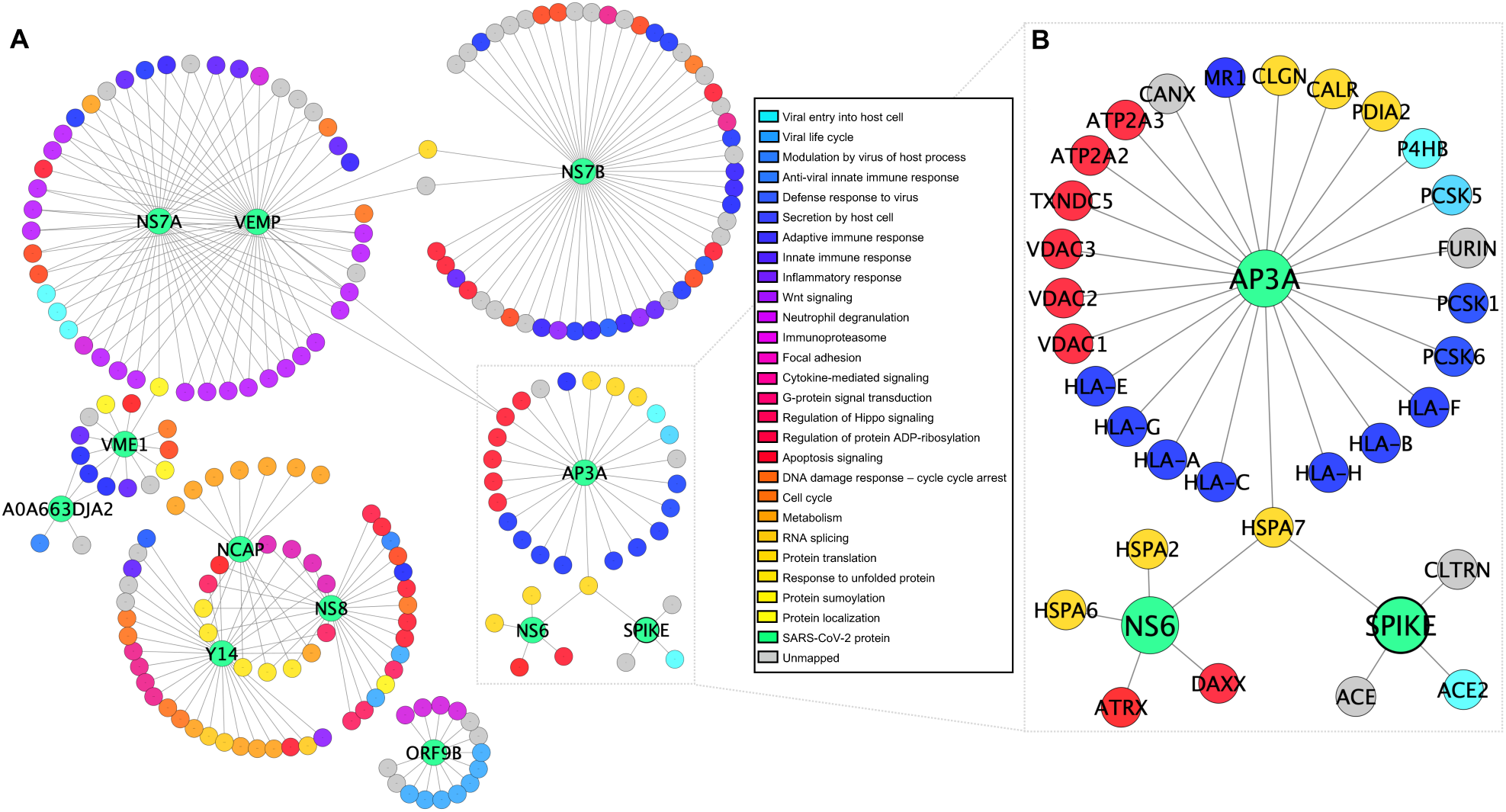
The Predicted SARS-CoV-2 vs. Human Interactome. GO-terms are approximately ordered by function, where related functions are closer in colour. The inset highlights the gene interactions of possibly greatest interest.

## 4 Conclusion

The purpose of this work is to help guide the broader research community in the collective pursuit to understand the SARS-CoV-2 viral pathogenesis. To that end, we assessed 285,124 protein pairs using two state-of-the-art sequence-based PPI predictors, thereby creating the comprehensive SARS-CoV-2 vs. human interactome. For each of the 14 SARS-CoV-2 proteins, a highly conservative locally defined decision threshold was determined to obtain a predicted interactome comprising 279 putative PPIs involving 225 human proteins within the predicted intersection of the PIPE4 and SPRINT methods. Furthermore, the PIPE-Sites algorithm was used to predict the putative interaction interfaces to identify the subsequence regions of interest that might mediate these interactions.

These predictions have been deposited in this public DataVerse repository for use by the broader scientific community in this collective effort to combat the COVID-19 pandemic [10]. All data and metadata are released under a CC-BY 4.0 licence and we re-emphasize that the information provided is theoretical modelling only and caution should be exercised in its use. It is intended only as a resource for the scientific community at large in furthering our understanding of SARS-CoV-2.

## Acknowledgements

This study was funded by a grant from the Canadian Natural Sciences and Engineering Research Council (NSERC) to JRG (RGPIN/327498-2011).

## Layperson Summary

The COVID-19 pandemic has led to catastrophic loss of life and the necessity to implement quarantines globally to contain the virus’ propagation. In order to reduce the transmission of the virus between people, three strategies can be used: wide-spread vaccination (only available in *∼*12-18 months), wide-spread use of anti-viral medication (available within months), and social distancing (immediately applicable, though highly disruptive). This paper focuses on the second strategy by using algorithms to study the novel coronavirus and to help the research community identify new ways to disrupt the virus.

We use the information available from other viruses that have infected humans (*e*.*g*. 2003 SARS virus, Dengue virus, Zika virus) to predict how physical interactions between human proteins and those from the novel coronavirus. We leveraged the Graham supercomputer to predict every possible relationship (>280,000) between nCoV and humans and generated these predictions. The algorithms are also used to identify the specific parts of the viral proteins that likely cause these interactions which is potentially useful to design drugs that can block that mechanism. This computational work is meant to function as a comprehensive “guide” to support other researchers in their exploration of new ways to protect against the novel coronavirus.

## Appendix

**Figure 10:**
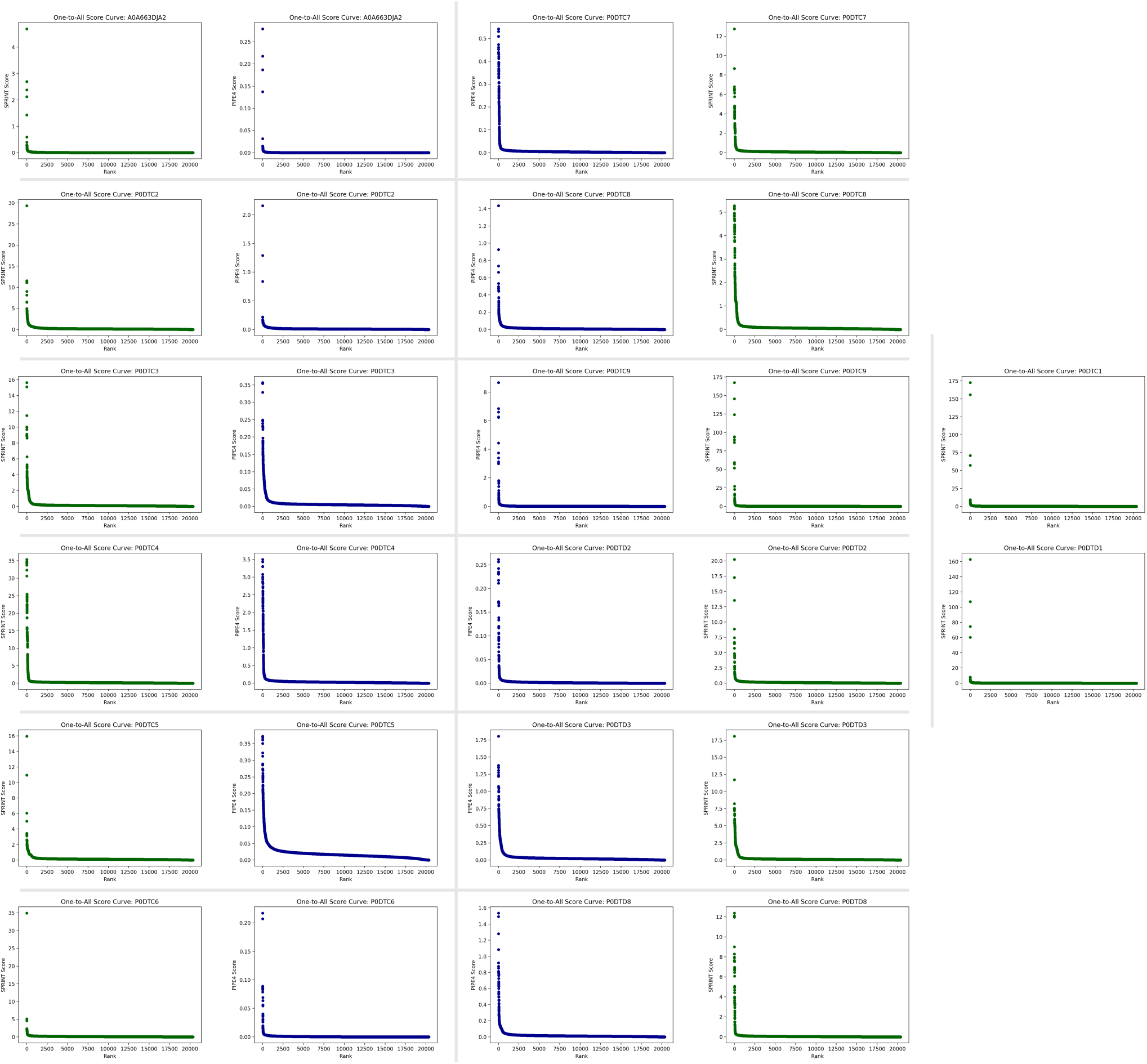
Compilation of the One-to-All Score Curves for each SARS-CoV-2 protein by PIPE4 (blue) and SPRINT (green). Each of the subplots depicts a characteristic “L”-shape, where there are a relatively small number of high-scoring pairs as compared to a large number of low-scoring pairs within the baseline. Note that the y-axes are not shared among subplots.

**Figure 11:**
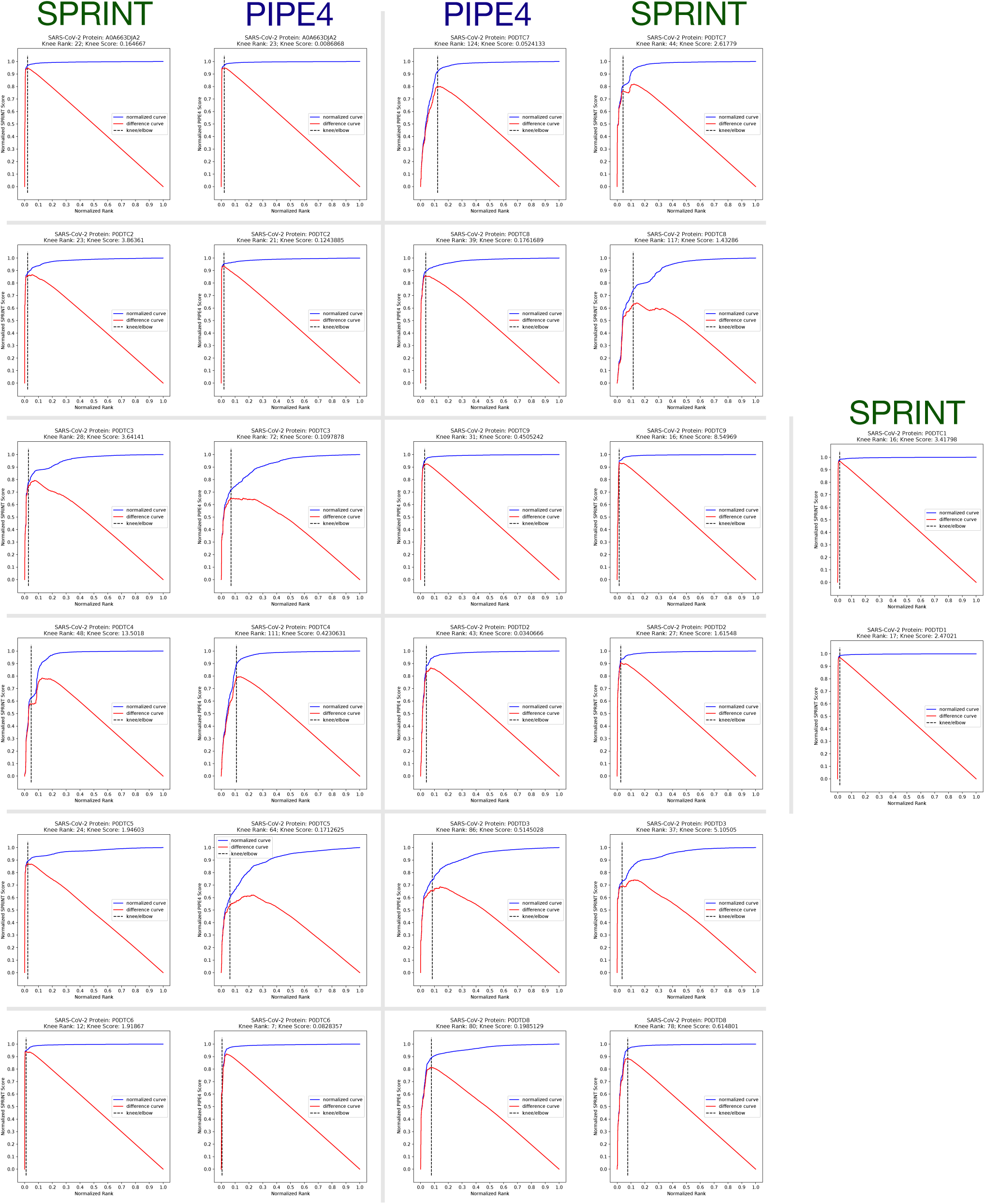
Compilation of the Detected Knee of each One-to-All Score Curves for each SARS-CoV-2 protein by PIPE4 and SPRINT. Each of the subplots highlights the detected knee of the normalized top-1000 predictions obtained using the Kneedle algorithm. The *differences* curve plots the value obtained from subtracting the perpendicular distance of each point to *y* = *x* from the distance of each point vertically to *y* = *x* of the normalized plot. The peak of this curve, parameterized by *S*, estimates the location of the knee.

**Figure 12:**
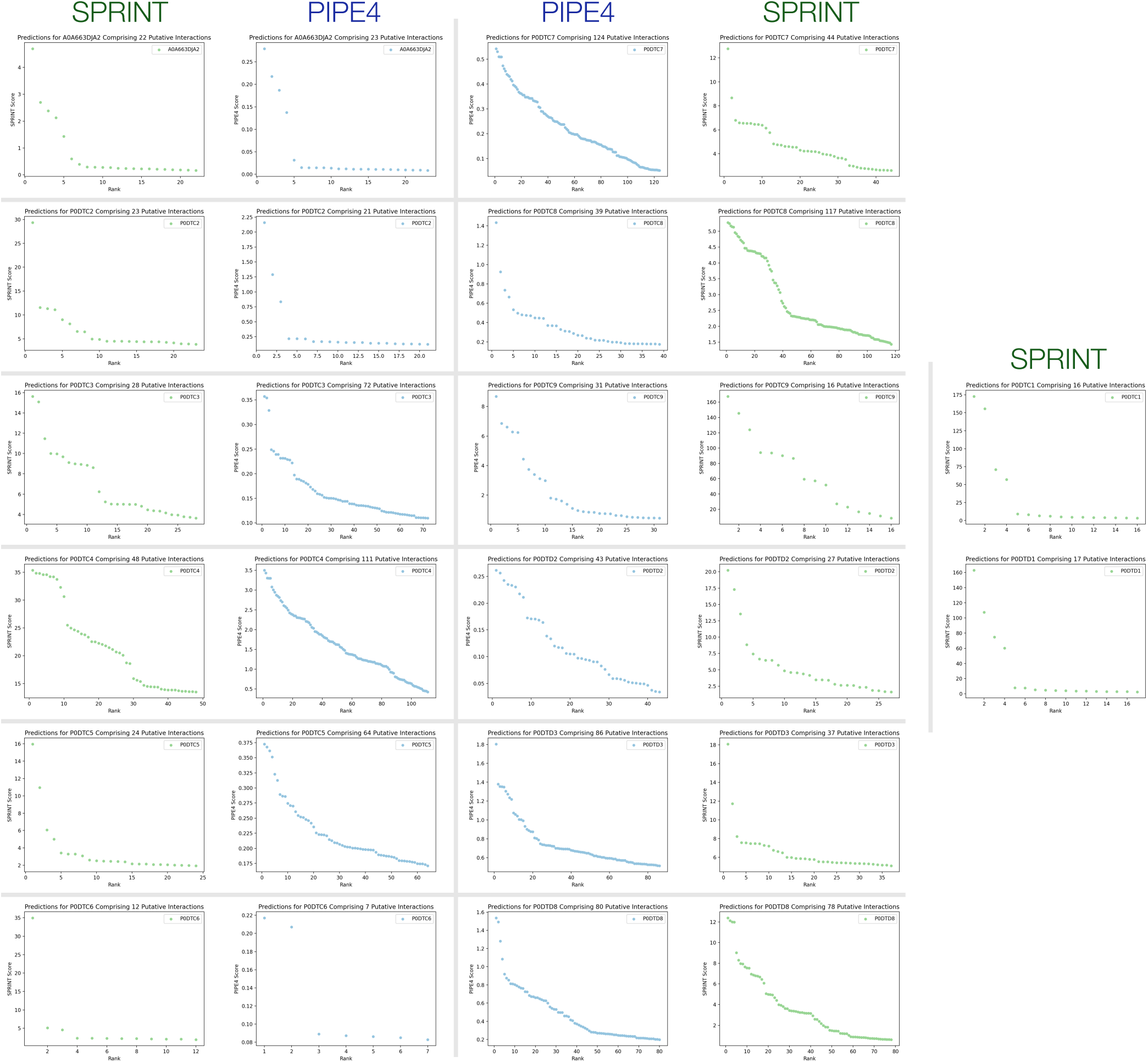
Compiled plot of all the Predicted Interactions for each protein and each method.

